# Altered Glia-Neuron Communication in Alzheimer’s Disease Affects WNT, p53, and NFkB Signaling Determined by snRNA-seq

**DOI:** 10.1101/2023.11.29.569304

**Authors:** Tabea M. Soelter, Timothy C. Howton, Amanda D. Clark, Vishal H. Oza, Brittany N. Lasseigne

## Abstract

**Background:** Alzheimer’s disease is the most common cause of dementia and is characterized by amyloid-β plaques, tau neurofibrillary tangles, and neuronal loss. Although neuronal loss is a primary hallmark of Alzheimer’s disease, it is known that non-neuronal cell populations are ultimately responsible for maintaining brain homeostasis and neuronal health through neuron-glia and glial cell crosstalk. Many signaling pathways have been proposed to be dysregulated in Alzheimer’s disease, including WNT, TGFβ, p53, mTOR, NFkB, and Pi3k/Akt signaling. Here, we predict altered cell-cell communication between glia and neurons.

**Methods:** Using public snRNA-sequencing data generated from postmortem human prefrontal cortex, we predicted altered cell-cell communication between glia (astrocytes, microglia, oligodendrocytes, and oligodendrocyte progenitor cells) and neurons (excitatory and inhibitory). We confirmed interactions in a second and third independent orthogonal dataset. We determined cell-type-specificity using Jaccard Similarity Index and investigated the downstream effects of altered interactions in inhibitory neurons through gene expression and transcription factor activity analyses of signaling mediators. Finally, we determined changes in pathway activity in inhibitory neurons.

**Results:** Cell-cell communication between glia and neurons is altered in Alzheimer’s disease in a cell-type-specific manner. As expected, ligands are more cell-type-specific than receptors and targets. We identified ligand-receptor pairs in three independent datasets and found involvement of the Alzheimer’s disease risk genes *APP* and *APOE* across datasets. Most of the signaling mediators of these interactions were not differentially expressed, however, the mediators that are also transcription factors had differential activity between AD and control. Namely, *MYC* and *TP53*, which are associated with WNT and p53 signaling, respectively, had decreased TF activity in Alzheimer’s disease, along with decreased WNT and p53 pathway activity in inhibitory neurons. Additionally, inhibitory neurons had both increased NFkB signaling pathway activity and increased TF activity of *NFIL3*, an NFkB signaling-associated transcription factor.

**Conclusions:** Cell-cell communication between glia and neurons in Alzheimer’s disease is altered in a cell-type-specific manner involving Alzheimer’s disease risk genes. Signaling mediators had altered transcription factor activity suggesting altered glia-neuron interactions may dysregulate signaling pathways including WNT, p53, and NFkB in inhibitory neurons.

## Background

Alzheimer’s disease (AD) is the most common cause of dementia, and an estimated 55 million people have dementia worldwide [1]. AD is characterized by amyloid-β plaques and phosphorylated tau tangles that result in neuronal loss, leading to memory deficits and overall cognitive decline [2]. While there are identified causes of familial early-onset AD, the causes of sporadic late-onset AD are largely unknown. Although neuronal loss is a primary hallmark of AD, non-neuronal cell populations are known to maintain brain homeostasis and neuronal health through neuron-glia and glial cell crosstalk via chemical messengers [3–6]. For example, astrocytes provide metabolic and nutritional support to neurons, while oligodendrocytes are responsible for neuron axon myelination [7–9]. Additionally, oligodendrocyte progenitor cells (OPCs) are in direct contact with neuronal synapses, and signaling between OPCs and other cell types, including glia and endothelial cells, has been previously indicated [10]. Microglia, as the resident macrophages of the central nervous system (CNS), release cytokines, chemokines, and growth factors in response to injury, which influence the synaptic activity of neurons [11–13]. Yet the glia-neuron interactions altered in AD have not been fully investigated despite being critical for understanding disease mechanisms and their potential as targets for therapeutic intervention. Between 2003 and 2021, no AD-mitigating therapeutics were approved by the FDA [14]. Aducanumab and lecanemab were approved in 2021 and 2023, respectively, despite concerns about their safety and efficacy, underscoring that the need for novel therapeutic targets persists.

Multiple signaling pathways have been proposed to be dysregulated in AD, including WNT (reviewed in [15, 16]), TGFβ (reviewed in [17–19]), mTOR (reviewed in [20, 21]), NFkB (reviewed in [22, 23]), p53 (reviewed in [24]), and Pi3k/Akt signaling (reviewed in [25–27]). Recently, multiple computational cell-cell communication (CCC) inference tools using single-nucleus RNA-sequencing (snRNA-seq) data to infer ligands and receptors were developed to predict signaling patterns between cell types (reviewed in [28]). Using these approaches, CCC patterns in the human postmortem prefrontal cortex (PFC) were previously shown to be disrupted in AD compared to control patients [29, 30]. For example, chandelier neurons were associated with an inhibitory signaling pattern in control PFC, though in AD they were associated with excitatory signaling involving the WNT signaling pathway [29]. Additionally, signaling involving glial cells, such as astrocytes and microglia, had an increased involvement of known AD-risk and neuroinflammatory genes in predicted ligands and receptors [29, 30]. While previous studies investigated overall patterns of altered CCC across all cell types without specifically describing altered glia-neuron interactions in-depth or focused on interactions between neurons and microglia [29–31], altered glia-neuron interactions and their downstream consequences in neurons remain understudied in AD.

Here, we used publicly available snRNA-seq AD and control data generated from postmortem human PFC to study altered glia-neuron interactions and their downstream effects in AD (**Fig 1A**) [32]. We inferred differential CCC interactions between astrocytes, microglia, oligodendrocytes, or OPCs (sender cell types) and inhibitory or excitatory neurons (receiver cell types; **Fig 1B**). We also investigated whether CCC is similar across cell types by calculating the Jaccard Similarity Index (JI) of ligands, receptors, and target genes between cell types (**Fig 1C**). We also validated our interactions using independent human PFC AD snRNA-seq datasets [33, 34] and further investigated the resulting high-confidence ligand-receptor pairs, their predicted downstream target genes, and signaling mediators through transcription factor (TF) and canonical signaling pathway activity (**Fig 1D-E**).

**Figure 1.**
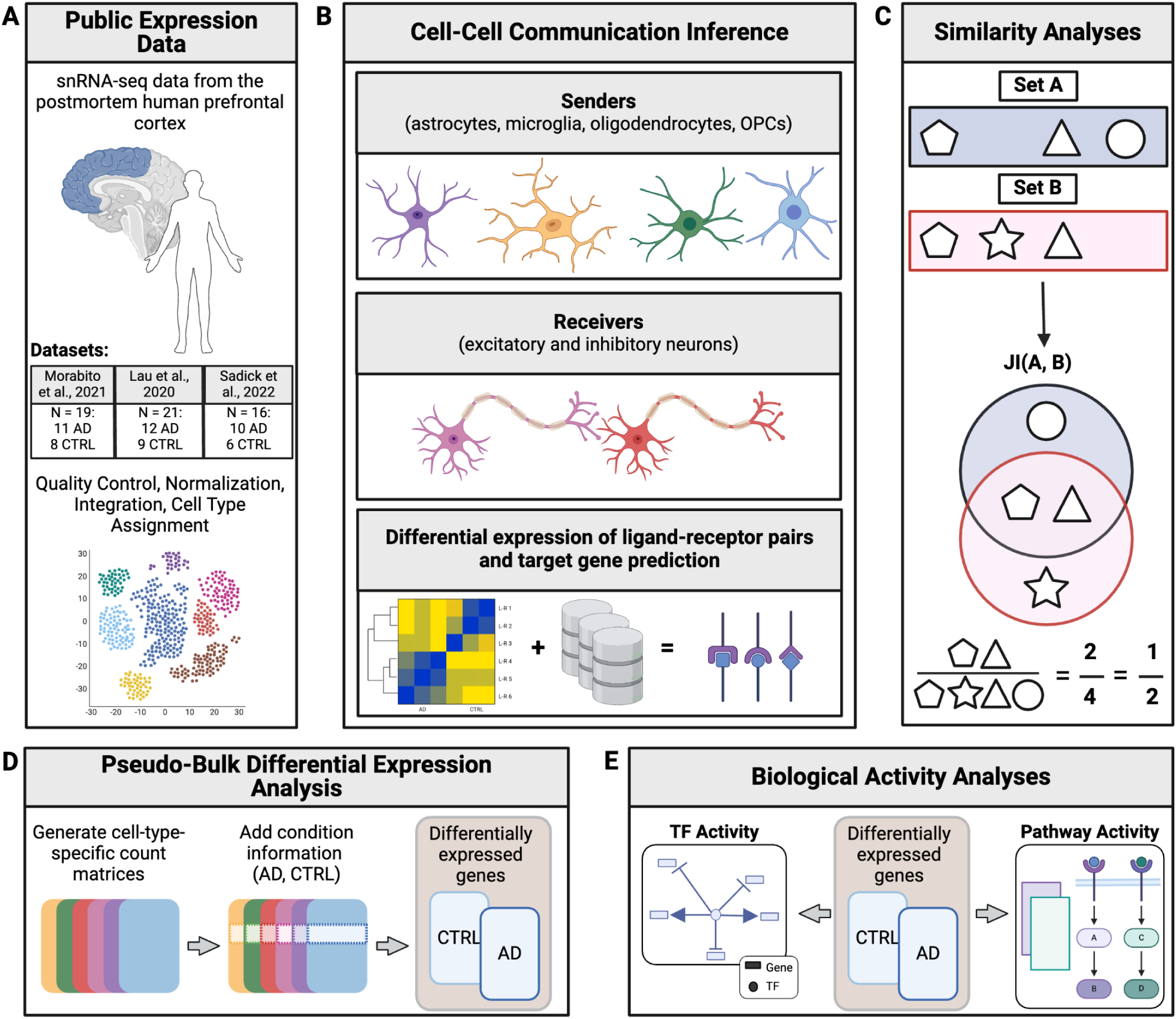
Schematic overview of our study design. **(A)** We downloaded and processed three publicly available snRNA-seq datasets from human postmortem prefrontal cortex (PFC) with Alzheimer’s disease (AD) as well as age- and sex-matched controls (CTRL). **(B)** We inferred ligand-receptor pairs between senders (astrocytes, microglia, oligodendrocytes, and oligodendrocyte progenitor cells - OPCs) and receivers (excitatory and inhibitory neurons) using gene expression and known ligand-receptor information. We also predicted target genes downstream of the ligand-receptor pairs [35]. **(C)** To determine whether interactions were cell-type-specific or shared, we calculated the Jaccard Similarity Index (JI) of ligands, receptors, and targets in senders and receivers. Shapes can refer to either ligands, receptors, or targets. **(D)** Then, we generated cell-type-specific count matrices to determine differentially expressed genes. **(E)** Using the differentially expressed genes, we inferred transcription factor (TF) activity of signaling mediators that are TFs and investigated canonical pathway activity [36].

## Methods

### Data Acquisition and Alignment

We downloaded fastq files of two publicly available snRNA-seq datasets (GSE174367 [32], GSE157827 [33]) generated from postmortem human PFC using the SRA-Toolkit v3.0.0. We also downloaded Cell Ranger processed files of one posrmotem human PFC snRNA-seq dataset (GSE167490 [34]) All datasets include AD as well as age- and sex-matched control samples (Morabito et al.: 11 AD and 8 control, Lau et al.: 12 AD and 9 control, Sadick et al.: 10 AD, 6 control). We aligned FASTQ files of the Morabito and Lau datasets to the 10x Genomics human reference genome (GRCh38) on the UAB Cheaha supercomputer using 10x Genomics Cell Ranger 6.1.1 [37].

### Data Processing and Quality Control

We performed all analyses in docker [38] with R v4.1.3 (**Availability of data and materials)**. Since Lau et al. had increased levels of ambient RNA after alignment, we performed ambient RNA removal using SoupX [39] v1.6.2 on Cell Ranger h5 files, which resulted in filtered Cell Ranger output matrices. According to 10x Genomics recommendation, we did not remove ambient RNA from Morabito et al., as there was no evidence of increased ambient RNA levels. Additionally, processed files of Sadick et al. did not include Cell Ranger h5 files, therefore we were not able to remove ambient RNA from this dataset. We used version 4.3.0.9002 of the Seurat R package for our analyses [40]. We combined Cell Ranger output matrices into sparse expression matrices using the *Read10x* function [40]. We created a Seurat object using the *CreateSeuratObject* function for each sample and condition (AD and control) before merging them into a single Seurat object. To evaluate data quality, we plotted bar plots of the total number of nuclei across conditions, density plots of the number of UMIs and genes, as well as the mitochondrial ratio and the number of genes per UMI, and correlation plots between the number of UMIs and genes colored by the mitochondrial ratio. We determined appropriate quality control cut-offs by visually inspecting these quality control plots for outliers or nuclei with high mitochondrial contamination. We performed quality control metrics at both the cell (i.e., mitochondrial ratio < 0.2, log10 of the number of genes per UMI > 0.8, and the number of genes per cell) and gene level (i.e., removing genes with non-zero counts in fewer than 10 cells to prevent skewing of average expression values) to control for the distributions of the quality control metrics and remove dead/dying cells. We filtered Morabito et al. and included cells with > 1,000 and < 10,000 genes per cell and we filtered Lau et al. to include cells with > 500 and < 10,000 genes per cell. We did not perform computational detection and removal of doublets, because doublet removal with DoubletFinder was not consistent or reproducible during internal peer-review of original code, yielding significant differences in the number of clusters and downstream cell type assignments. Then, using default parameters, we performed batch correction within each dataset using harmony v0.1.0 [41], as it preserves biological variation while reducing variation due to technical noise [42]. We also performed Principal Component Analysis (PCA) using the *RunPCA* function from Seurat [40] without approximation (approx = FALSE) for complete reproducibility. We scaled and normalized features/genes before plotting UMAPs to confirm successful integration across conditions.

### Clustering and Cell Type Identification

In order to determine clustering resolution, we compared multiple resolutions between 0 and 2 using clustree v0.5.0 for each dataset [43]. Since we did not remove doublets, we focused on identifying clustering resolutions with the fewest switches of nuclei between clusters, as doublets can have hybrid transcriptomes of multiple cell types [44]. We chose resolutions 1.2, 1, and 0.9, which identified 32, 34, and 25 clusters using leiden v0.4.3 [45] for Morabito et al., Lau et al., and Sadick et al., respectively. We identified differentially expressed marker genes for each cluster (using the *FindAllMarkers* function on the RNA assay from Seurat [40]) with a log fold change threshold > 0.2 and a significant Bonferroni adjusted p-value. We assigned cell types using differential expression of cell-type-specific genes (**Tables S1 and S2**) identified through PanglaoDB [46], CellMarker [47], and the Human Protein Atlas [48] at https://www.proteinatlas.org/ and feature plots using canonical cell type markers (**Table S3**). We sub-clustered neuron clusters initially unidentifiable as excitatory or inhibitory using the *FindSubCluster* function from Seurat [40] and assigned their cell type through differential expression (log fold change threshold > 0.25) of marker genes. Due to extensive mixing of marker gene expression indicative of doublets, we removed two unknown clusters (clusters 20 and 25) in Sadick et al.

### Cell-Cell Communication Inference

To infer CCC between glia and neurons across conditions (AD and control), we applied multinichenetr v1.0.0 and used the Nichenet v2 prior [35]. We converted our processed Seurat objects to Single Cell Experiment objects using the SingleCellExperiment R package [49] v1.16.0 before using them as input to MultiNicheNet. Then, we performed ligand-receptor pair prediction between all cell types in each dataset and filtered for senders (astrocytes, microglia, oligodendrocytes, and OPCs) and receivers (excitatory and inhibitory neurons) between conditions. Since MultiNicheNet uses pseudobulk aggregation for its differential expression analysis, we only included samples with a minimum of 5 cells per cell type per sample. We identified differentially expressed ligands, receptors, and target genes using a 0.5 logFC threshold with non-adjusted p-values of 0.05, as recommended by the package developers. We performed principal component analysis (PCA) and looked at multiple clinical covariates and based on distinct separation and clustering due to sex, we included sex as a covariate for Morabito et al. and Lau et al. (PCA; **Fig S1 & S2**). When calculating ligand activity, we considered the top 250 targets with the highest regulatory potential. We prioritized ligand-receptor interactions by min-max scaling the scores for all comparisons of the following metrics: differential expression of ligand, differential expression of the receptor, the fraction of ligand-receptor pairs expressed across samples, expression of ligand, expression of the receptor, scaled ligand activity, the abundance of the sender in the condition of interest, and abundance of the receiver in the condition of interest. Then, we performed weighted aggregation of the scaled scores using default MultiNicheNet prioritization weights. We calculated expression correlation information between ligand-receptor pairs and their targets by calculating Pearson and Spearman correlation coefficients. Downstream analyses only included ligand-receptor-target (LRT) interactions with Spearman and Pearson correlation coefficients greater than 0.33 and less than −0.33, as recommended. Finally, we compiled a list of high-confidence LRTs. High-confidence interactions were LRTs that met previous filtering criteria in the Morabito and Lau datasets while involving the same sender and receiver cell type.

### Similarity Calculations of Ligands, Receptors, and Target Genes Across Cell Types

We calculated the JI to determine the degree of overlap of ligands, receptors, and target genes between sender or receiver cell types. First, we calculated JI between receiver cell types (excitatory and inhibitory neurons) based on the overlap of receptors and target genes. Then, we calculated JI between sender cell types (astrocytes, microglia, oligodendrocytes, and OPCs) based on the overlap of ligands, receptors, and target genes.

### Gene Regulatory Network Inference

For every high-confidence LRT between Morabito et al. and Lau et al., we generated gene regulatory networks (active signaling networks) that included the top 2 regulators (based on whether they were upstream of the target and downstream of the ligand) in each dataset with the *get_ligand_signaling_path_with_receptor* function from multinichenetr v1.0.0 R package [35]. We scored and ranked signaling regulators based on whether they were upstream of the target gene and downstream of the ligand. We assigned edge weights to the networks by combining them with MultiNicheNet’s prior knowledge, which includes information on ligand-receptor to target signaling paths. Then, we investigated the active signaling networks’ topology using the igraph v1.5.1 R package [50]. We generated weighted and directed igraph objects for individual LRT-specific active signaling networks using the *graph_from_data_frame* function. Then, we determined all potential signaling mediators by identifying nodes outgoing from the receptor node using the *neighbors* function. Finally, we determined the shortest path between the receptor and target gene nodes using the *shortest_paths* function using Dijkstra’s algorithm to infer the most likely path and, therefore, the top mediator(s) of signal transduction of our high-confidence interactions (**Table 1**).

**Table 1.**
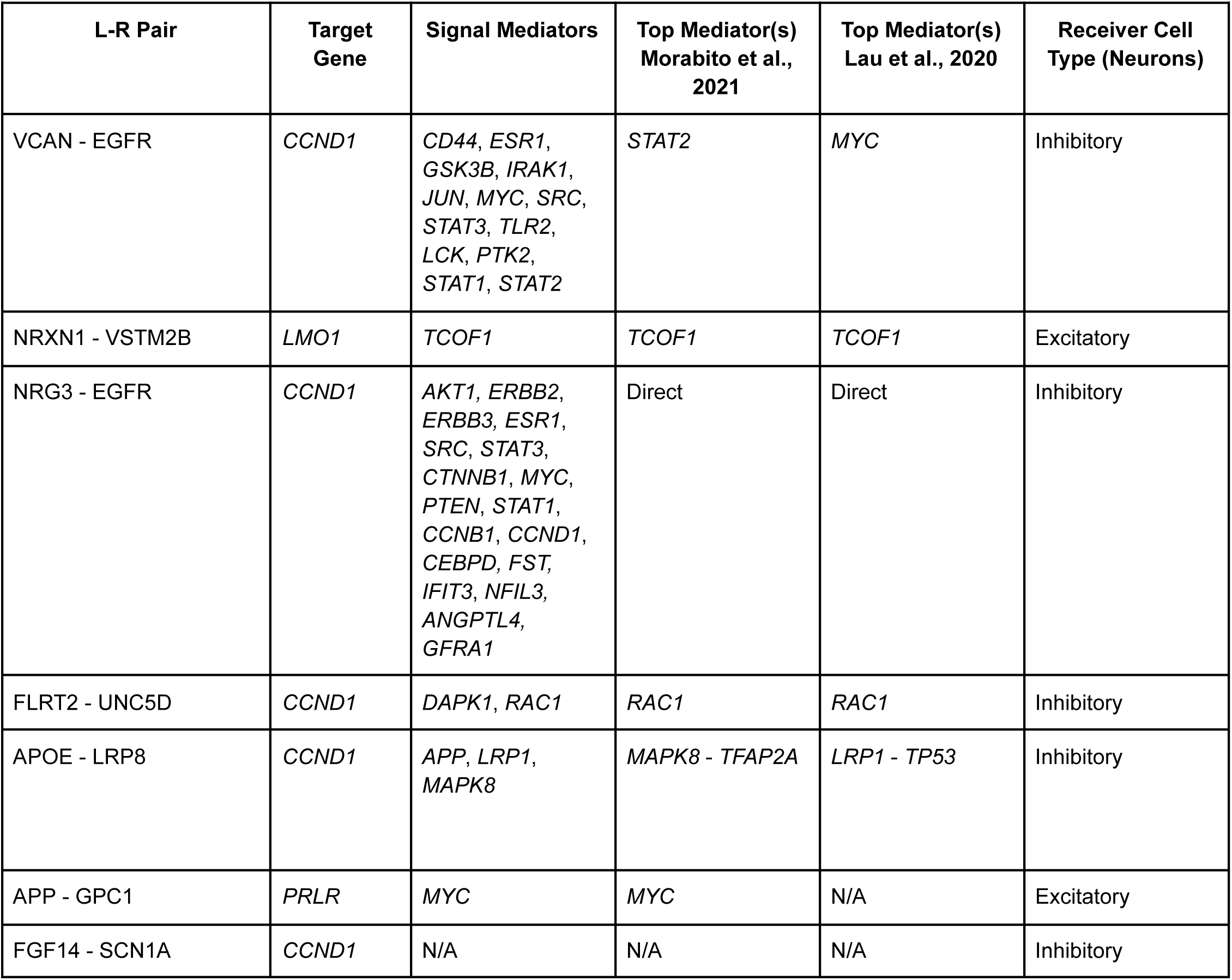
Signaling mediators of ligand-receptor pairs and their downstream targets across datasets.

| L-R Pair | Target Gene | Signal Mediators | Top Mediator(s)<br>Morabito et al.,<br>2021 | Top Mediator(s)<br>Lau et al., 2020 | Receiver Cell<br>Type (Neurons) |
| --- | --- | --- | --- | --- | --- |
| VCAN - EGFR | *CCND1* | *CD44, ESR1, GSK3B, IRAK1, JUN, MYC, SRC, STAT3, TLR2, LCK, PTK2, STAT1, STAT2* | *STAT2* | *MYC* | Inhibitory |
| NRXN1 - VSTM2B | *LMO1* | *TCOF1* | *TCOF1* | *TCOF1* | Excitatory |
| NRG3 - EGFR | *CCND1* | *AKT1, ERBB2, ERBB3, ESR1, SRC, STAT3, CTNNB1, MYC, PTEN, STAT1, CCNB1, CCND1, CEBPD, FST, IFIT3, NFIL3, ANGPTL4, GFRA1* | Direct | Direct | Inhibitory |
| FLRT2 - UNC5D | *CCND1* | *DAPK1, RAC1* | *RAC1* | *RAC1* | Inhibitory |
| APOE - LRP8 | *CCND1* | *APP, LRP1, MAPK8* | *MAPK8 - TFAP2A* | *LRP1 - TP53* | Inhibitory |
| APP - GPC1 | *PRLR* | *MYC* | *MYC* | N/A | Excitatory |
| FGF14 - SCN1A | *CCND1* | N/A | N/A | N/A | Inhibitory |

### Pseudo-bulking of snRNA-seq Data

To overcome the data sparsity in snRNA-seq [51], we pseudo-bulked the Morabito et al. and Lau et al. datasets and generated cell-type-specific count matrices, as previously described [52]. We converted raw data from the Seurat object *counts* slot into Single Cell Experiment objects using SingleCellExperiment [49] v1.16.0 while also incorporating necessary metadata with information on sample origin (sample_id), condition (group_id), and cell type (cluster_id). We aggregated counts for each cell type across samples using the *aggregate.Matrix* function (Matrix.utils R package v0.9.7), yielding gene by sample counts for each cell type. We appended the metadata with condition information (AD or control) in preparation for our differential gene expression analysis (see below).

### Differential Gene Expression Analysis

We performed differential gene expression analysis between conditions (AD vs control) using DESeq2 v1.34.0 for each cell type [53] in Morabito et al. and Lau et al. First, we created dds objects using the *DESeqDataSetFromMatrix* function. We made pairwise comparisons for every cell type between conditions using the Wald test, while including sex as a covariate, as determined by PCA (**Fig S1 & S2**). Then, we performed log fold change shrinkage to generate more accurate log2foldchange estimates. While DESeq2 recommends the application of Bayesian log fold change shrinkage (type = apeglm), we used its original shrinkage estimator (type = normal), as it preserves the Wald test statistic (*stat*) needed for our downstream analyses of TF and pathway activity.

### Transcription Factor Activity Analysis

To determine whether transcription factors (TFs) that are signaling mediators had activator or repressor roles in inhibitory neurons, we used the Wald test statistic (*stat*) from previously generated pseudo-bulk differential gene expression analysis outputs. To obtain one dataframe for the Morabito and Lau datasets, we combined *stat* scores from both datasets into one data frame before inferring TF activity. However, as datasets did not have the same number of identified differentially expressed genes and we did not aim to only investigate genes shared between datasets, we replaced NAs with 0, because decoupleR cannot interpret NAs. We used the CollecTRI prior (accessed in September 2023), which has information on TFs, including their direction of regulation on their targets [54], and combined it with pseudo-bulked count matrices for inhibitory neurons. We calculated activity scores for all TFs in inhibitory neurons that had a minimum of 5 targets using the Multivariate Linear Model (*run_mlm* function decoupleR v2.7.1 [36]), where the t-values from the model represent regulator activity [36]. Positive values represent an activator role, while negative values represent a repressor role in AD compared to control. After inferring TF activity, we subsetted for TFs that were signaling mediators in our high-confidence LRTs in inhibitory neurons.

### Pathway Activity Analyses

Similarly to our TF activity analysis, we used the Wald test statistic (*stat*) from pseudo-bulk differential gene expression analysis outputs to infer pathway activity in inhibitory neurons. As before, we replaced NAs with 0, as the datasets did not have the same number of differentially expressed genes and decoupleR is unable to interpret NAs. We used the top 500 responsive genes ranked by p-value in PROGENy, a collection of pathways and their target genes (accessed in October 2023) [55]. We calculated pathway activity scores for all pathways with a minimum of 5 targets using the Multivariate Linear Model (*run_mlm* function decoupleR v2.7.1 [36]). Positive values represent an increase, while negative values represent a decrease in pathway activity in AD compared to control.

## Results

### Identification of cell-cell communication patterns from AD snRNA-seq data

To infer cell-cell communication (CCC) patterns in AD, we used previously published snRNA-seq data generated from 19 postmortem human PFC, including 11 AD and 8 age- and sex-matched control samples (Morabito et al., 2021, **Fig 2**) [32]. We aligned raw FASTQ files and retained ∼65K nuclei after quality control. We identified 32 distinct clusters using Leiden clustering [45], and assigned eight major brain cell types using differential expression of canonical marker genes (**Fig 2A-B, Tables S1-3**). Cell types were evenly distributed and integrated across conditions (**Fig. 2C**). Next, we determined whether cell types were balanced across conditions. We observed that all cell type proportions were slightly biased towards AD (**Fig 2D).** However, as there is an imbalance of AD to control samples in the dataset (11 and 8, respectively), we expected this.

**Figure 2.**
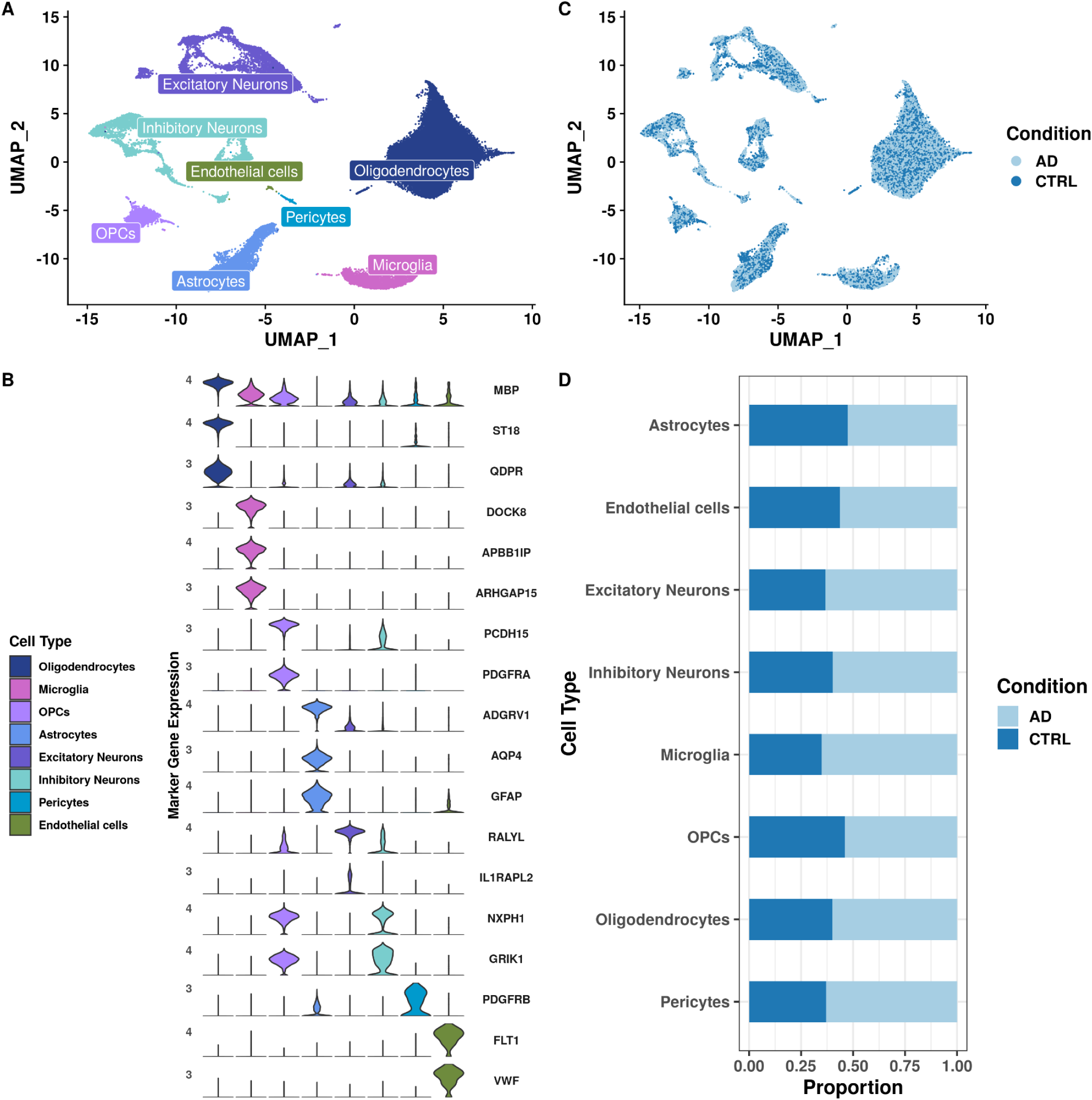
Overview of our approach and dataset. **(A)** Uniform Manifold Approximation and Projection (UMAP) of 65,546 nuclei colored by their assigned cell types. **(B)** Violin plot of canonical marker gene expression used for cell type assignment. Color represents cell type. **(C)** UMAP of integrated dataset split by condition (AD, CTRL). **(D)** Stacked barplot of cell type proportions across conditions (AD, CTRL).

### Ligand, receptor, and target gene patterns are mostly cell-type-specific in AD

We predicted interactions using MultiNicheNet [35] by determining genes differentially expressed between AD and control in all cell types before filtering for our sender (astrocytes, microglia, oligodendrocytes, and OPCs) and receiver (excitatory and inhibitory neurons) cells of interest (**Fig 1B**). By cross-referencing differentially expressed genes with the NicheNet prior, we identified more than 45,000 differentially expressed ligand-receptor interactions between AD and control across all cell types (**Fig S3A**). After filtering interactions based on our senders and receivers of interest (**Fig S3B**) and prioritizing interactions based on ligand activity and regulatory potential,[35] we were left with 1,766 ligand-receptor-target (LRT) pairings (**Fig 3A**). Out of 1,766 predicted interactions, 1,519 were up-regulated in AD and 247 were up-regulated in control, suggesting their decreased expression and, therefore a loss of signal in AD (**Fig S3C**). Most interactions originated from OPCs (n = 704), followed by astrocytes (n = 442), oligodendrocytes (n = 318), and microglia (n = 302). Interestingly, OPCs had the fewest (n = 3,097), and oligodendrocytes had the most nuclei (n = 38,864). In contrast, astrocytes and microglia had 5,272 and 3,862 nuclei, respectively, suggesting that the number of predicted interactions is independent of the number of nuclei in each cell type. Of the 1,766 prioritized LRTs, 801 were specific to excitatory, and 965 were specific to inhibitory neurons (**Fig 3A**). To determine whether CCC patterns were cell-type-specific or shared, we compared ligands, receptors, and target genes by JI. JI scores range from 0 (no overlap, i.e., cell-type specific) to 1 (total overlap, i.e., shared). In order to determine the similarity between excitatory and inhibitory neuron signaling patterns, we calculated their JI with respect to senders for receptor and target genes only. Since ligands are specific to senders and not receivers, we did not evaluate their overlap in excitatory and inhibitory neurons. Overall, receptors and target genes were not highly similar between neuronal subtypes (JI < 0.5 for 7/8 comparisons; **Fig 3B**). Interestingly, we observed overall higher JI in receptors (JI 0.18 - 0.53) than in target genes (JI 0.09 - 0.15), indicating differential downstream effects despite overlapping receptors between excitatory and inhibitory neurons (**Fig 3B**). In addition to comparisons between receiver cell types, we were interested in the similarity of ligands, receptors, and target genes between sender cell types by neuron subtypes (**Fig 3C-H**). For ligands associated with inhibitory neurons, the greatest overlap was between ligands from OPCs and microglia (JI = 0.09), closely followed by oligodendrocytes and OPCs (JI = 0.07) interactions (**Fig 3C**). For ligands interacting with excitatory neurons, oligodendrocytes had the highest similarity with astrocytes (JI = 0.14; all other comparisons JI < 0.12; **Fig 3F**). Receptor JI scores were higher overall than ligands, and Microglia and OPCs had the highest similarity score with astrocytes for inhibitory and excitatory neurons (JI = 0.22, **Fig 3D** and JI = 0.19, **Fig 3G**, respectively). Finally, the target genes of inhibitory and excitatory neurons had a higher JI than ligands or receptors (**Fig 3E & 3H**). The greatest overlap of target genes in inhibitory neurons was between microglia and astrocytes (JI = 0.45; **Fig 3E**), despite microglia- and astrocyte-expressed ligands having low similarity (JI = 0.05; **Fig 3C**). Within inhibitory neurons, OPCs and microglia shared the least number of target genes (JI = 0.32; **Fig 3E**). We observed the same pattern of high target gene similarity with low ligand similarity between senders in excitatory neurons (e.g., microglia-astrocytes: target JI = 0.33 and ligand JI = 0.04, **Fig 3F and 3H**). Overall, OPCs and astrocytes showed similar and consistent overlap across receptors and targets with both inhibitory and excitatory neurons (**Fig 3C-H**). Then, we performed the same analyses in an independent snRNA-seq dataset (Lau et al., 2020) [33] from the same brain region and with similar sequencing depth and cell type composition. The Lau et al. dataset included 21 human postmortem PFC samples (12 AD, 9 control). As with the Morabito et al. data, we aligned FASTQ files and assigned cell types using differential expression of canonical marker genes (**Fig S4A-B**). We integrated ∼32,000 cells across conditions (**Fig S4C**), which resulted in 7 cell types (**Fig S4A**). Finally, the cell type proportions were balanced across conditions for all but the endothelial cells, which had an increased proportion in AD (75% AD, 25% control; **Fig S4D**). After differential CCC analysis, we identified fewer total interactions (n = 969) in the Lau et al. data, and unlike before in Morabito et al., we predicted less interactions specific to inhibitory neurons (n = 464) compared to excitatory neurons (n = 505; **Fig S5A**). Similarly, receptors and target genes were not highly similar between neuronal subtypes (JI < 0.12 for all comparisons, **Fig S5B**), and target genes between senders had a higher JI than ligands and receptors in both excitatory and inhibitory neurons in this validation dataset (**Fig S5C-H**). Generally, the results from our independent validation dataset recapitulate our original findings. Overall, we found that ligands, receptors, and target genes are largely cell-type-specific between and across sender and receiver cell types in these two datasets. This supports cell-type-specific altered CCC patterns in AD.

**Figure 3.**
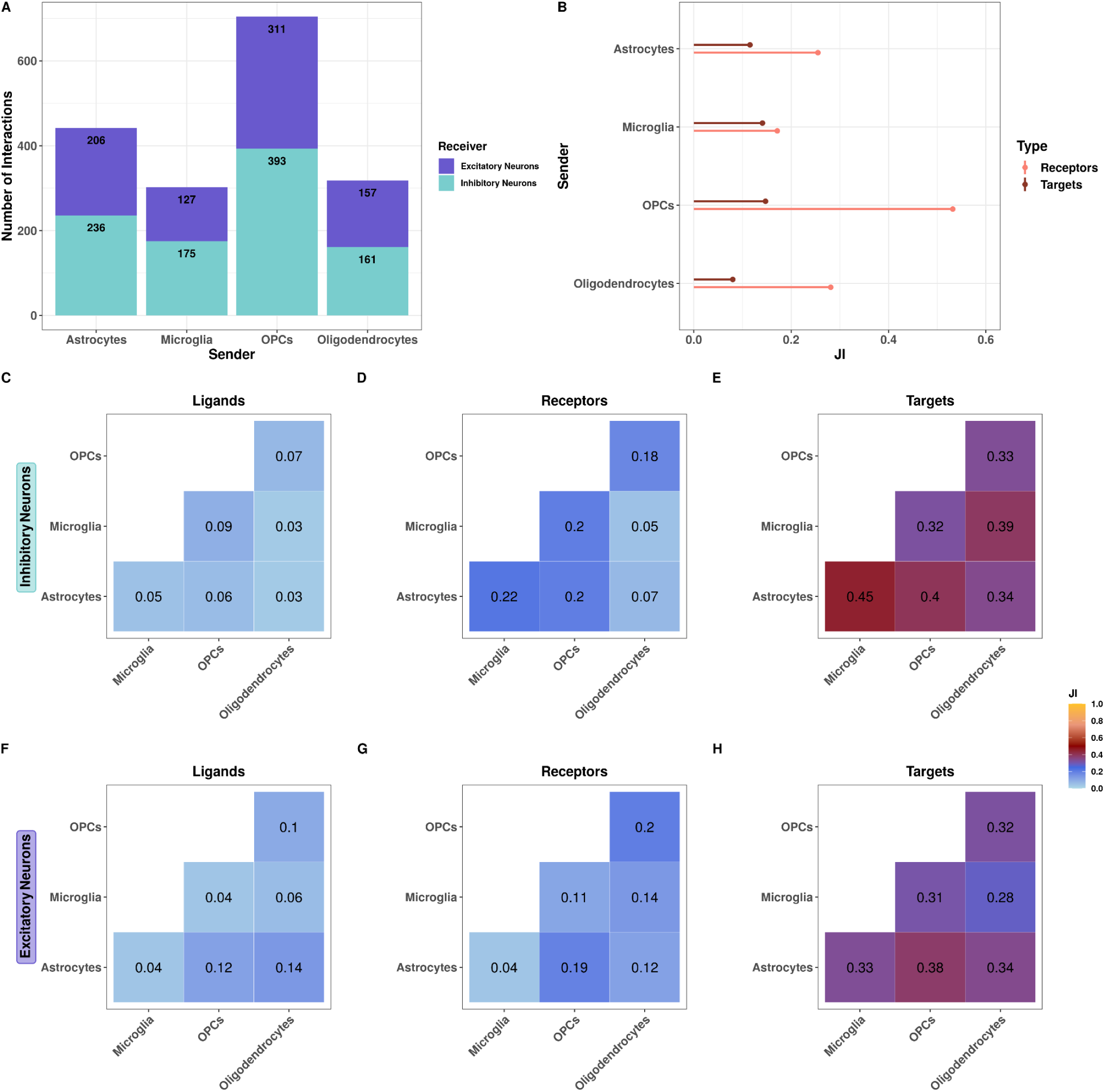
Ligand, receptor, and gene target patterns are mostly cell-type-specific in AD. **(A)** Stacked barplot of the number of prioritized interactions where excitatory or inhibitory neurons are the receiver. **(B)** Jaccard Similarity Index (JI) between excitatory and inhibitory neurons of receptors and targets. JI between sender cell type **(C)** ligands, **(D)** receptors, and **(E)** targets for inhibitory neurons. JI between sender cell type **(F)** ligands, **(G)** receptors, and **(H)** targets for excitatory neurons.

### Analyses of altered CCC in 3 datasets reveal shared ligand-receptor pairs and downstream target genes

To determine high-confidence interactions that were conserved across multiple AD patient cohorts, we determined the overlap of altered CCC interactions (both ligand-receptor and LRTs) identified in three datasets. In addition to the Morabito and Lau datasets, we included a third postmortem human PFC snRNA-seq dataset (Sadick et al.) with more than 100k nuclei from 10 AD and 6 control samples (**Fig S6A-B;** [34]). We predicted 20 ligand-receptor pairs that were shared between all three independent patient cohorts, which included an interaction with the AD-risk gene *APP* (**Fig S7A & S7C**). The interaction with *APP* as a ligand was also the only LRT we confirmed across all datasets (**Fig S7B**). We excluded the Sadick et al dataset from downstream analyses as it had the broadest range of patient age (56 - 100 years), the fewest samples (total n = 16), and was enriched specifically for astrocytes and oligodendrocytes, which resulted in neuronal nuclei depletion [34]. Additionally, the cell type proportions were imbalanced across condtions (possibly due to enrichment of specific glia; **Fig S6C-D**), which affects downstream analyses [56]. We also had no information on the average sequencing depth, while the Morabito and Lau datasets had comparable sequencing depths (per sample average mean reads per cell > 95k for both datasets).

When we compared Morabito and Lau, we identified 46 ligand-receptor pairs that were upregulated in AD with the same sender and receiver cell type across datasets, comprised of 29 ligands and 32 receptors (**Fig S8A**). Out of 29 ligands, only FLRT2 was predicted in more than 1 cell type (**Fig S8**). Additionally, we identified 10 ligand-receptor pairs that were upregulated in control, which we inferred as interactions lost or downregulated in AD. Interestingly, these interactions included 2 AD-risk genes as ligands: APP and APOE, predicted in oligodendrocytes and astrocytes, respectively (**Fig S8B**). Except for OPCs, we find that ligand-receptor pairs overlapping across datasets seem to be receiver-specific, as few interactions are shared between excitatory and inhibitory neurons in astrocytes, microglia, and oligodendrocytes (**Fig 4A**). Additionally, oligodendrocytes were the only sender across both datasets that had overlapping communication with excitatory neurons only (**Fig 4A**). We also predicted APP as a receptor in excitatory neurons originating from OPCs (**Fig 4A**). Additionally, we also identified APP as a ligand in oligodendrocytes interacting with GPC1 in excitatory neurons, while the risk factor APOE functions as a ligand in astrocytes interacting with LRP8 in inhibitory neurons (**Fig 4A**). Both of these interactions were downregulated in AD. We identified EGFR as an inhibitory neuron receptor in 3 of 4 senders (oligodendrocytes had no overlapping interactions with inhibitory neurons; **Fig 4A**), which has been proposed as a therapeutic target in AD [57, 58]. However, ligands targeting EGFR were dependent on the sender cell type. Out of 56 ligand-receptor pairs identified in both datasets, only 7 also shared the same downstream target gene (**Fig 4B**). In comparison, the 49 remaining ligand-receptor pairs did not show target gene overlap (**Table S4**). Our 7 high-confidence LRTs included interactions originating from three of four senders (microglia had no overlapping interactions) and converged to three target genes: *LMO1*, *PRLR*, and *CCND1* (**Fig 4B**). Interestingly, most of our high-confidence interactions originated from OPCs and astrocytes (n = 3 each). The interactions involving the AD-risk genes *APP* and *APOE,* as well as *EGFR,* also remained among our high-confidence interactions. Additionally, pseudo-bulk expression information for the three overlapping target genes confirms the same gene expression directionality across datasets in AD, where *LMO1* and *PRLR* were upregulated in excitatory neurons, while *CCND1* was downregulated in inhibitory neurons across datasets (**Fig 4C**). Finally, AD is associated with sex effects [59] and indeed, removing sex as a covariate in our models identifed 17 LRTs (**Fig S9;** 10 not identified with sex as a covariate), suggesting there may be potential sex differences in altered glia-neuron communication in AD. Altogether, we showed that ligand-receptor pairs are mostly receiver-specific and that the majority of our senders, except for oligodendrocytes, had altered CCC with both excitatory and inhibitory neurons across AD datasets. Our findings also indicate that interactions involving AD-risk or AD-associated genes (like *EGFR*) are preserved across different patient cohorts.

**Figure 4.**
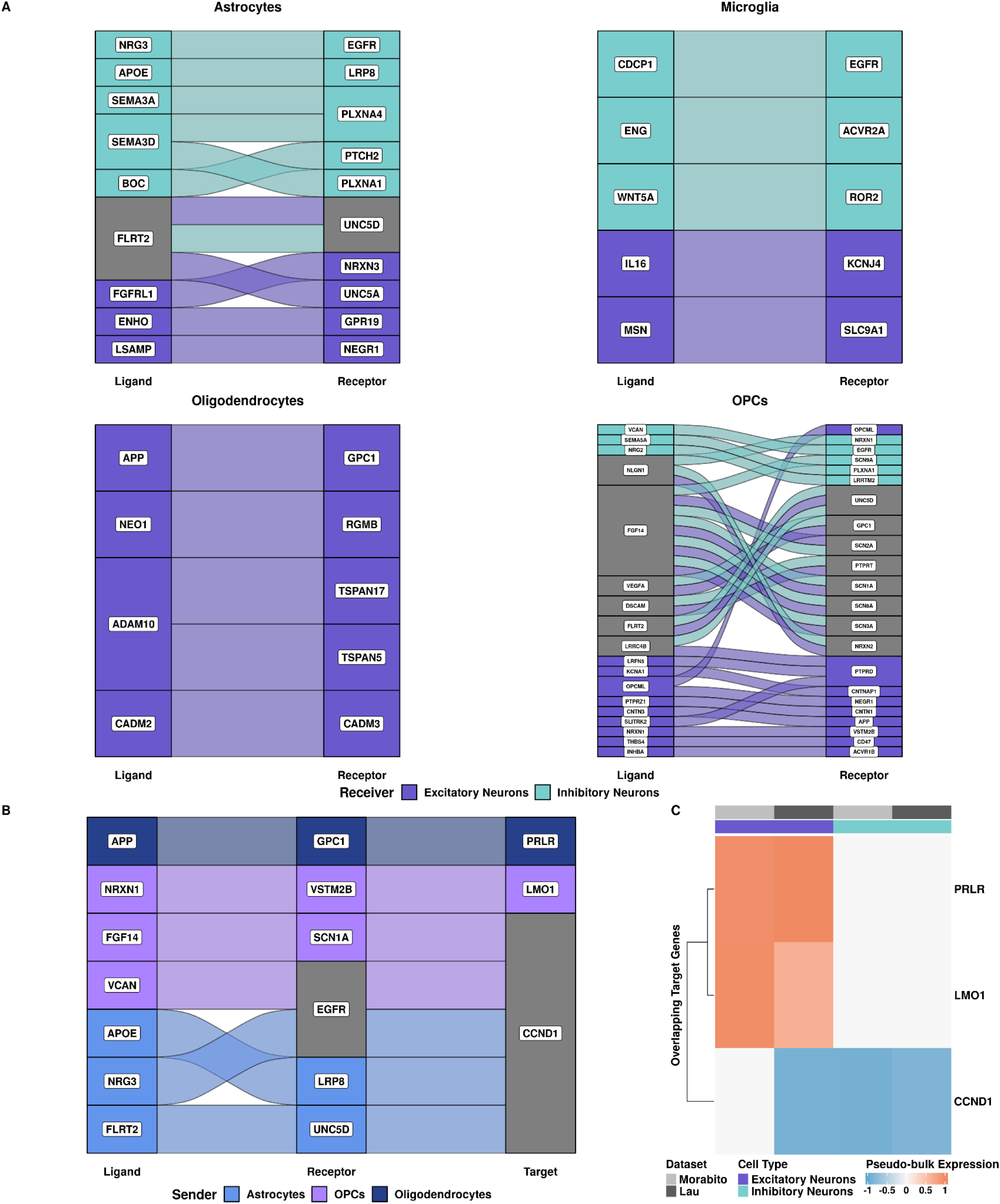
Ligand-receptor pairs and target genes overlap between 2 independent snRNA-seq datasets from the same brain region. **(A)** Alluvial plots of overlapping ligand-receptor pairs for each sender cell type colored by receiver cell type. **(B)** Alluvial plot of the 7 ligand-receptor-target (LRT) pairings that overlap across datasets, colored by sender cell type. Grey indicates the involvement of more than one sender cell type. **(C)** Heatmap of pseudo-bulk expression values for target genes which were identified in both datasets annotated by receiver cell type and dataset.

### Signaling mediators are not differentially expressed but show altered TF and pathway activity in inhibitory neurons

We constructed gene regulatory networks for all high-confidence interactions using ligand-receptor to target signaling information from MultiNichNet [35] to identify signaling mediators downstream of predicted ligand-receptor pairs. In total, we identified 32 signaling mediators across datasets in inhibitory neurons (**Table 1**). We considered any node adjacent to our receptor node as a potential signaling mediator and predicted the most likely signaling modulators using shortest path calculations with Dijkstra’s algorithm between predicted receptors and targets. First, we performed a differential gene expression analysis on pseudo-bulked count data from inhibitory neurons to investigate whether signaling modulators were differentially expressed in AD compared to control. Interestingly, 16 of 32 signaling mediators had divergent gene expression across our datasets, and we observed that only *CCND1* was significantly downregulated in both (**Fig 5A**). Additionally, two genes were significantly different in AD in the Morabito et al. data (*ANGPTL4* and *CD44*; p-value < 0.05), however, neither *ANGPTL4* nor *CD44* was significant in the Lau et al. data. Since most of our predicted signaling mediators were not differentially expressed in AD, this could indicate that their downstream effects may be due to gene regulatory mechanisms instead of gene expression, therefore, we investigated TF activity. Using pseudo-bulked gene expression information from inhibitory neurons, we calculated TF activity scores for 10 signaling mediators that were also TFs. While none of the TFs were significantly differentially expressed, 5 out of 10 had divergent gene expression patterns across datasets (gene expression was increased in only one dataset and vice versa, **Fig 5A**). Overall, MYC, CTNNB1, STAT2, and TP53 consistently function as repressors, suggesting that they downregulate their target genes in AD inhibitory neurons across datasets (**Fig 5B**). MYC, a cancer-associated cell cycle regulator [60], showed the greatest overall significant difference in TF activity in AD (**Fig 5B**). *MYC* had slightly increased expression (logFC of 0.015 and 0.103 in Morabito et al. and Lau et al., respectively; **Fig 5A**) while downregulating its targets in AD (**Fig 5B**). Additionally, CTNNB1, which is associated with cell-cell adhesion regulation and coordination [61], also significantly downregulates its targets in AD in the Morabito et al. dataset (**Fig 5B**). Like *MYC*, *CTNNB1* also had increased expression in AD (logFC of 0.023 and 0.025 in Morabito et al. and Lau et al., respectively; **Fig 5A**). Both *CTNNB1* and *MYC* are involved in the WNT signaling pathway [62], which was decreased in AD (**Fig 5C**). Interestingly, despite its divergent gene expression pattern (downregulated in Morabito et al., upregulated in Lau et al.; **Fig 5A**), TP53 functions as atranscriptional repressor in AD (**Fig 5B**), which is further recapitulated by decreased p53 signaling pathway activity in inhibitory neurons (**Fig 5C**). NFIL3, associated with NF-kappaB (NFkB)[63], was the only TF to function as transcriptional activators across both datasets in AD (**Fig 5B**). Along with high TF activity, we observed increased *NFIL3* expression (**Fig 5A**) and activity of the NFkB signaling pathway (**Fig 5C**) in AD, which were both consistent across datasets. All other TFs, including CEBPD, JUN, ESR1, STAT1, and STAT3 showed divergent TF activity across datasets. STAT1, STAT3, and ESR1 activity was increased in Lau et al. and decreased in Morabito et al., while JUN and CEBPD activity was reversed (**Fig 5B**). Interestingly, STAT1 and STAT3 TF activities were different across datasets (**Fig 5B**); therefore it was not surprising that JAK-STAT signaling in AD was decreased in Morabito et al. and increased in Lau et al. (**Fig 5C**). Overall, we find consistently altered TF and pathway activity associated with WNT, p53, and NFkB signaling dysregulation in inhibitory neurons across datasets despite the lack of significant gene expression changes of signaling mediators. This further suggests gene regulatory consequences of altered CCC between glia and neurons in AD.

**Figure 5.**
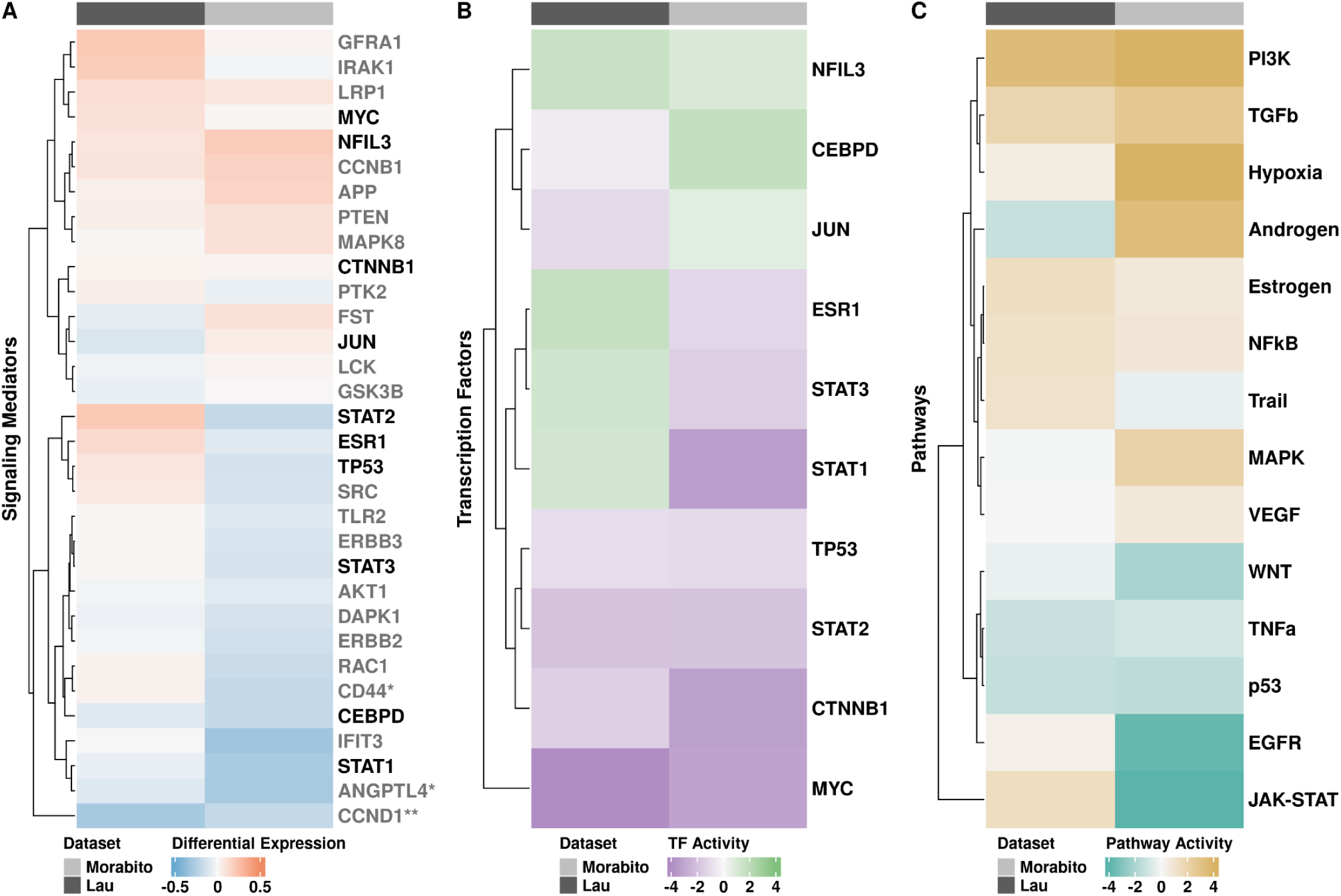
Signaling mediators in inhibitory neurons are not differentially expressed but show altered TF and pathway activity in AD. **(A)** Heatmap representing pseudo-bulk differential gene expression of potential signaling mediators in inhibitory neurons annotated by dataset. Asterisks indicate significant differential gene expression (** in both datasets and * in Morabito et al.). Mediators that are TFs are denoted in black. Red and blue represent negative and positive logFC values. **(B)** Heatmap of TF activity scores that are different between AD and control of signaling mediators, annotated by dataset. Green and purple represent increased and decreased TF activity, respectively. **(C)** Heatmap of pathway activity scores of all differentially expressed genes in AD from DESeq2 annotated by dataset. Tan and teal represent pathway activity as increased or decreased, respectively.

## Discussion

In this study, we investigated altered CCC in publicly available snRNA-seq datasets from postmortem human PFC in AD using *in silico* approaches to generate high-confidence LRT interactions. Previous *in silico* studies have either investigated overall CCC patterns across all brain cell types without specifically describing glia-neuron interactions and their downstream effects in depth or focused on interactions between excitatory neurons and microglia [29–31]. Here, we predicted interactions between glia (astrocytes, microglia, oligodendrocytes, and OPCs) and neurons (excitatory and inhibitory), while also investigating their potential downstream consequences in inhibitory neurons. Although neuronal loss is a primary hallmark of AD, glia are important for maintaining brain homeostasis and neuronal health through interactions with neuronal cell populations [3–6]. For example, astrocytes provide metabolic and nutritional support [7, 8], oligodendrocytes myelinate neuronal axons [9], microglia respond to injury affecting neuronal function [11–13], and OPCs make direct contact with neuronal synapses [10]. Consistent with previous studies [29, 30], our JI analyses of ligands, receptors, and targets confirmed that interactions between glia and neurons are altered in a cell-type-specific manner in AD. Although ligands are known to target many receptors across different cells, causing a variety of downstream effects [64, 65], a previous study in hematopoietic stem cells revealed greater cell-type-specificity in ligands than receptors [66]. Our findings corroborate this in the PFC, as we observed the greatest cell-type-specificity in ligands, followed by receptors and then targets. Finally, since our high-confidence interactions converge to 3 downstream target genes, it is plausible that although ligand-receptor pairs are sender-specific, the aggregated effects of altered glia-neuron communication converge in both excitatory and inhibitory neurons.

Our high-confidence interactions included *APP*, a familial AD risk gene, and *APOE*, the strongest genetic risk factor of sporadic AD [67], which has previously been confirmed as a ligand for another AD-risk gene, *TREM2*. A recent study has shown the involvement of APOE as a ligand in astrocyte-neuron interactions through the receptors LRP1 and SORL1 in both inhibitory and excitatory neurons [31]. Interestingly, we predicted an interaction of APOE with a different low-density lipoprotein receptor, LRP8, between astrocytes and inhibitory neurons. Even though the association of LRP8 with AD was not robust in previous work [68], the APOE-LRP8 interaction was among our high-confidence interactions. Additionally, low-density lipoprotein receptors have high and isoform-specific binding affinity for *APOE* [69]. Combined, this suggests a potential role for receptors in the low-density lipoprotein receptor family in AD that requires further investigation.

Semaphorins are signaling proteins that most often bind to plexin receptors crucial to neurodevelopment and the adult CNS [70]. Immunohistochemistry work in the human brain illustrated that semaphorin3A (SEMA3A) colocalizes with tau neurofibrillary tangles in neurons during the later stages of AD [71]. SEMA3A has also been indicated to induce activation of cyclin-dependent kinase 5 (CDK5), promoting phosphorylation of tau neurofibrillary tangles [71, 72]. Our data further support these initial findings, as we predicted 2 ligand-receptor pairs involving semaphorin ligands and plexin receptors across three independent datasets (SEMA3A - PLXNA4 and SEMA3D - PLXNA4). However, the predicted target genes of these ligand-receptor pairs were different across datasets. Addtionally, our initial analyses without patient sex as a covariate indicate that interactions between glia and neurons, including semaphorin-plexin interactions that target *SMAD7,* are potentially sex-biased. Therefore, performing our analyses both before and after accounting for patient sex led us to conclude that there are potential sex differences in altered glia-neuron communication in AD. As almost two thirds of patients are women [73], AD has a prominent female sex bias, that we are able to capture at the CCC level. Thus, sex biases in altered CCC in AD should be further investigated. Additionally, SEMA6D interacts with TREM2 through altered neuron-microglia interactions, regulating microglia activation in AD [30]. This further indicates the importance of semaphorins in AD, as their signaling is altered between multiple sender and receiver cell types.

Canonical signaling through the WNT [15, 16], p53 [24], and NFkB [22, 23] signaling pathways have been shown to be dysregulated in AD. The WNT signaling pathway is crucial for normal brain function and neuronal survival in the adult CNS and has been extensively implicated in AD (reviewed in [15, 16]). Interestingly, our pathway activity analyses indicated decreased WNT signaling in AD inhibitory neurons. Increased levels of Wnt signaling have been shown to have a protective role against amyloid-β plaques in both *in vitro* and *in vivo* models of AD [74–76]. We also identified *MYC* and *CTNNB1* as potential signaling mediators of high-confidence interactions in inhibitory neurons. *MYC* is at the crossroads of multiple signaling pathways, including WNT signaling [62], and its increased expression leads to elevated vulnerability of neurons in degenerative diseases [77]. *CTNNB1* is a key player in the WNT signaling pathway [61]. In concordance with the literature, we observed increased expression of *MYC* and *CTNNB1*, increased repressor TF activity for both, and overall decreased WNT signaling in AD. Furthermore, the p53 signaling pathway is often associated with carcinogenesis due to its role in DNA damage and cellular stress responses, but there is mounting evidence of its involvement in AD. *In vitro* data indicates increased levels of p53 expression [78], which we also observed. Additionally, the inactivation of the p53 pathway is known to lead to cell cycle re-entry of senescent cells [79], and, in AD, neuronal cell cycle re-entry exacerbates neuronal loss [80, 81]. While we found different gene expression patterns of *TP53* across datasets, we identified altered TF activity of TP53 and decreased p53 signaling pathway activity in AD inhibitory neurons consistent across datasets. The NFkB signaling pathway regulates the expression of proinflammatory genes in the context of immunity [82]. It has also been shown to be associated with neuroinflammation in AD (reviewed in [22, 23]), as well as amyloid-β plaque and tau neurofibrillary tangle pathologies (reviewed in [83]). Multiple studies have shown that amyloid-β accumulation in AD leads to neurotoxic activation of the NFkB signaling pathway [83], which may lead to activation of Beta-site APP cleaving enzyme 1 (BACE1), initiating the formation of amyloid-beta through splicing of *APP* [84]. Interestingly, we observed differential *NFIL3* gene expression between AD and control and activator TF activity in AD of NFIL3, an important regulator of the NFkB signaling pathway, consistent across datasets. We also observed increased activity of the NFkB signaling pathway in inhibitory neurons in AD. Altogether, this suggests that altered glia-neuron interactions may play a role in dysregulating canonical signaling pathways like WNT, p53, and NFkB signaling in inhibitory neurons in AD.

Although our study provides insight into dysregulated glia-neuron interactions and their downstream effects in AD, there are limitations including the use of curated priors for CCC and biological activity inference and postmortem data with varying sequencing depth and number of nuclei. Our CCC inference relies on curated priors, therefore, any interactions that have not been described previously are excluded from our prior, limiting our ability to predict entirely novel interactions. Additionally, we inferred protein abundances of ligands and receptors from gene expression information. Due to multiple processes like post-translational modifications and protein degradation, mRNA and protein levels are not always directly related. Therefore, our hypothesis-generating approach calls for further confirmation of predicted interaction through additional experimentation (e.g.,RNAscope or co-culture) in future studies with access to appropriate samples. Similarly, TF and pathway activity analyses also rely on previously curated priors. Finally, despite our best efforts to choose comparable datasets, there was extensive technical variability between datasets (number of nuclei and sequencing depth). While AD is a heterogeneous and multifactorial disorder, we expected greater LRT overlap. However, despite the little LRT overlap across datasets, which may be due to the technical variability between snRNA-seq datasets or the gene regulatory information in the MultiNicheNet prior, there was a consistent overlap of predicted ligand-receptor pairs across multiple independent patient cohorts. These ligand-receptor pairs should be further investigated for their specific role and potential druggability in AD. Additionally, as more snRNA-seq datasets with increased sequencing depth and number of nuclei become available, our analyses should be expanded to confirm and extend findings from the present study. Finally, we are using data generated from postmortem human brains, a time point where extensive neuronal degeneration and loss has occurred. Therefore, we are likely not capturing the altered CCC patterns relevant to disease etiology and progression. Future studies should investigate altered signaling across time points or disease stages (e.g., in AD mouse models) to properly address how disease progression affects CCC in AD.

## Conclusions

Overall, using public snRNA-seq data from postmortem human PFC, we inferred altered cell-type-specific CCC between glia and neurons in AD. We find that the AD-risk genes *APP* and *APOE* are among altered interactions conserved across patient cohorts. Additionally, we observed altered TF activity of signaling mediators, along with altered signaling pathway activity, in inhibitory neurons. Therefore, our findings suggest that altered glia-neuron interactions may dysregulate canonical signaling in pathways like WNT, p53, and NFkB through TFs and co-regulators like MYC, CTNNB1, TP53, EP300, and NFIL3.

## Abbreviations

AD: Alzheimer’s disease
OPC: oligodendrocyte progenitor cell
CNS: central nervous system
CCC: cell-cell communication
snRNA: seq single-nucleus RNA sequencing
PFC: prefrontal cortex
JI: Jaccard Similarity Index
TF: transcription factor
CTRL: control
PCA: Principal Component Analysis
UMAP: Uniform Manifold Approximation and Projection
LRT: ligand-receptor-target
SEMA3A: semaphorin3A
CDK5: cyclin-dependent kinase 5

## Declarations

## Ethics approval and consent to participate

Not applicable

## Consent for publication

Not applicable

## Availability of data and materials

The snRNA-sequencing datasets used in this study were downloaded from the GEO database (GSE174367 [32], GSE157827 [33], GSE167490 [34]). The data of this study are available on Zenodo (doi: 10.5281/zenodo.10214497). The code supporting the results of this study is available on Zenodo (doi: 10.5281/zenodo.10211622) and GitHub (https://github.com/lasseignelab/230313_TS_CCCinHumanAD). Docker images with R version 4.1.3 used for these analyses are publicly available on Zenodo (doi: 10.5281/zenodo.10214660) and Docker Hub (https://hub.docker.com/repository/docker/tsoelter/rstudio_ccc_ad/general).

## Authors’ contributions

TMS, VHO, TCH, and BNL conceptualized the project. TMS downloaded and performed preprocessing. TMS adapted code from VHO for Jaccard Similarity Index calculations. All other analyses were coded and performed by TMS. TCH, ADC, and VHO reviewed and validated the code. BNL provided supervision and project administration. BNL and TMS acquired funding. TMS wrote the first draft. TMS, TCH, ADC, VHO, and BNL reviewed and edited the manuscript. All authors read and approved the final manuscript.

## Acknowledgments

The authors thank the Lasseigne Lab members Jordan Whitlock, Emma Jones, and Elizabeth Wilk for their feedback throughout this study. We also thank the UAB Biological Data Science group (RRID:SCR_021766) for providing a script for helping to run containers on the UAB high-performance cluster (https://github.com/U-BDS/training_guides/blob/main/run_rstudio_singularity.sh).

