## Supplemental Information for "Altered Glia-Neuron Communication in Alzheimer’s Disease Affects WNT, p53, and NFkB Signaling Determined by snRNA-seq"

**Figure S1: PCA plots before and after correction for patient sex in the Morabito et al. data.** PCA plots colored by sex for all cell types of interest before adjustment for sex (left column) and after inclusion as a covariate (right column).

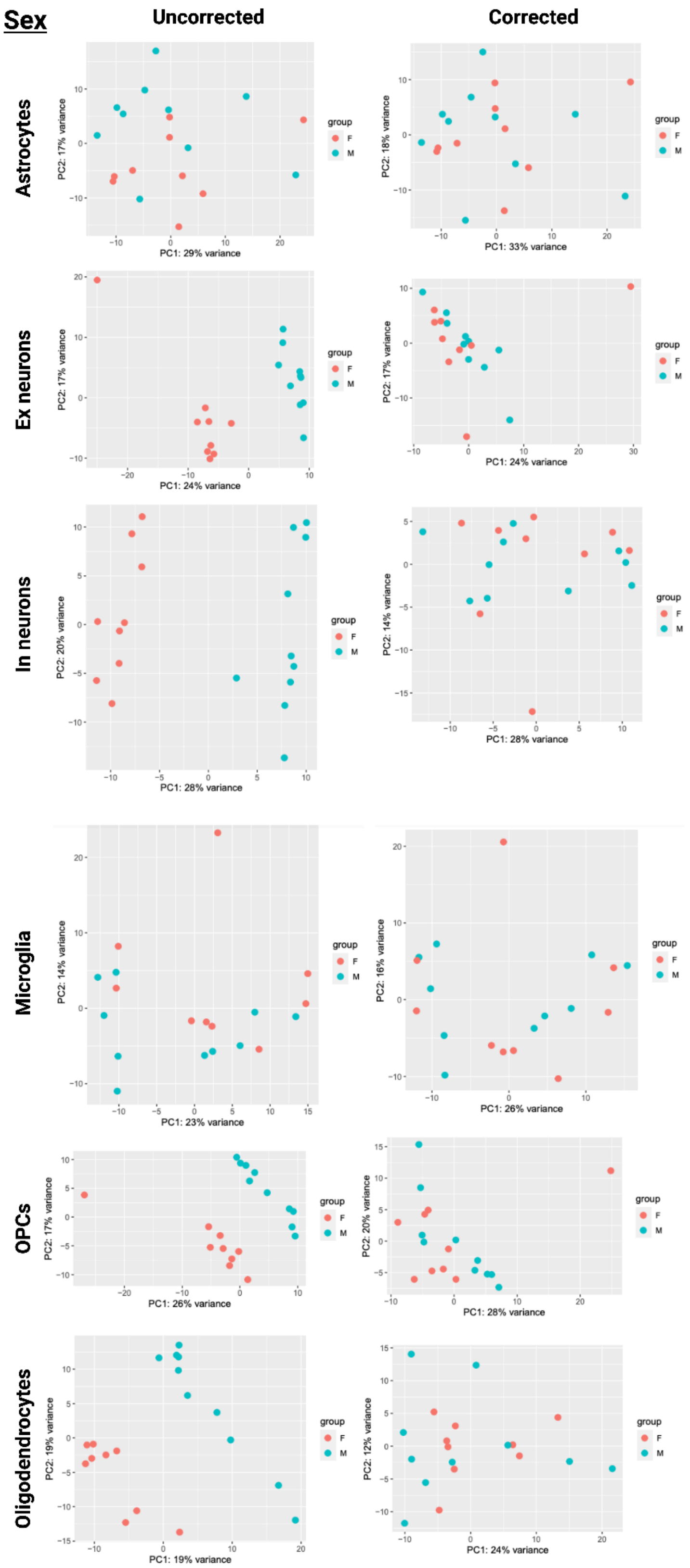

**Figure S2: PCA plots before and after correction for patient sex in the Lau et al. data.** PCA plots colored by sex for all cell types of interest before adjustment for sex (left column) and after inclusion as a covariate (right column).

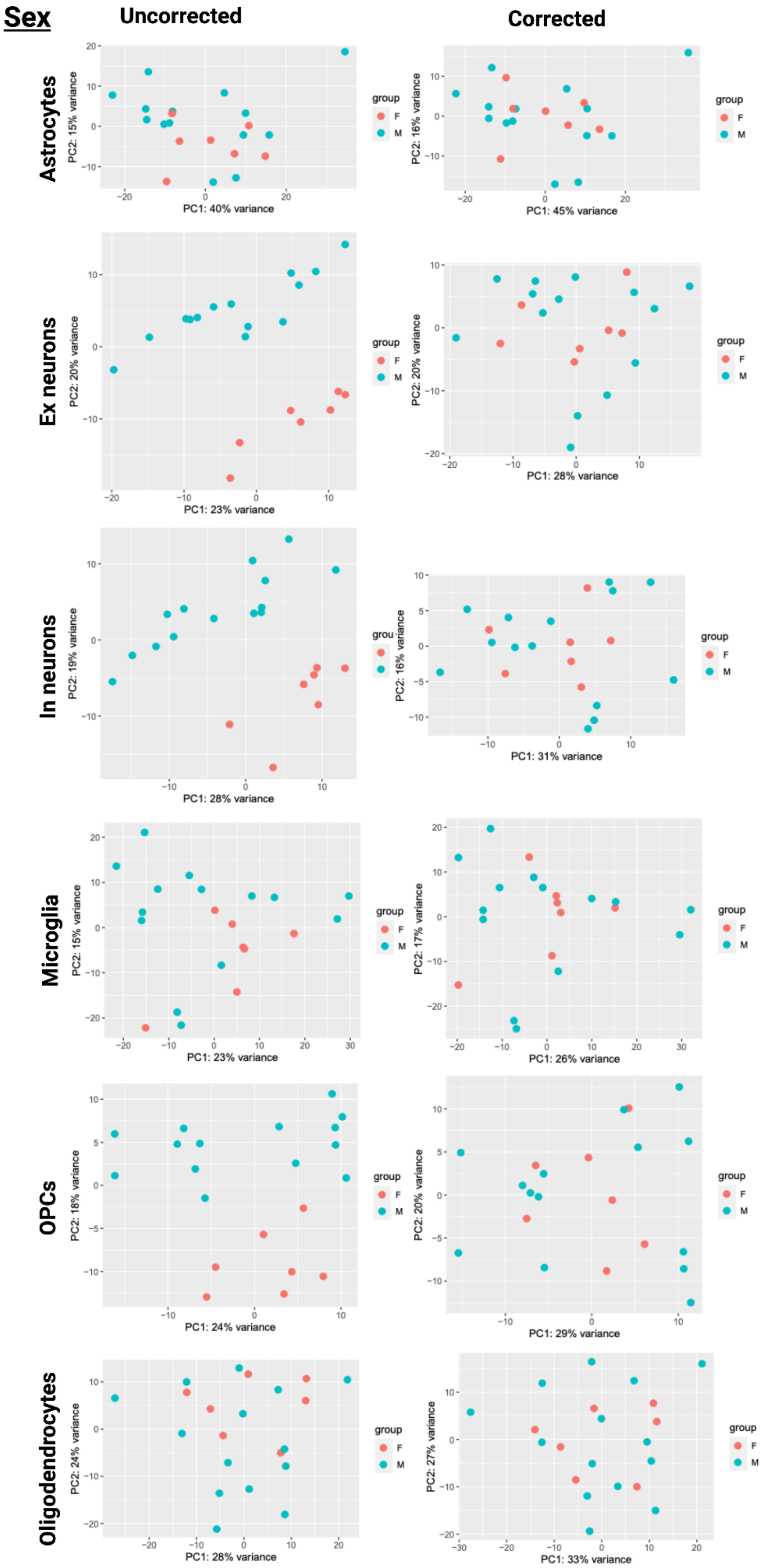

**Figure S3: Number of cell-cell communication interactions.**

(**A**) Barplot showing the total number of predicted interactions across all cell types in AD. (**B**) Total number of interactions filtered by sender and receiver (excitatory and inhibitory neurons) cell types of interest, and (**C**) after prioritization using ligand activity and regulatory potential scores.

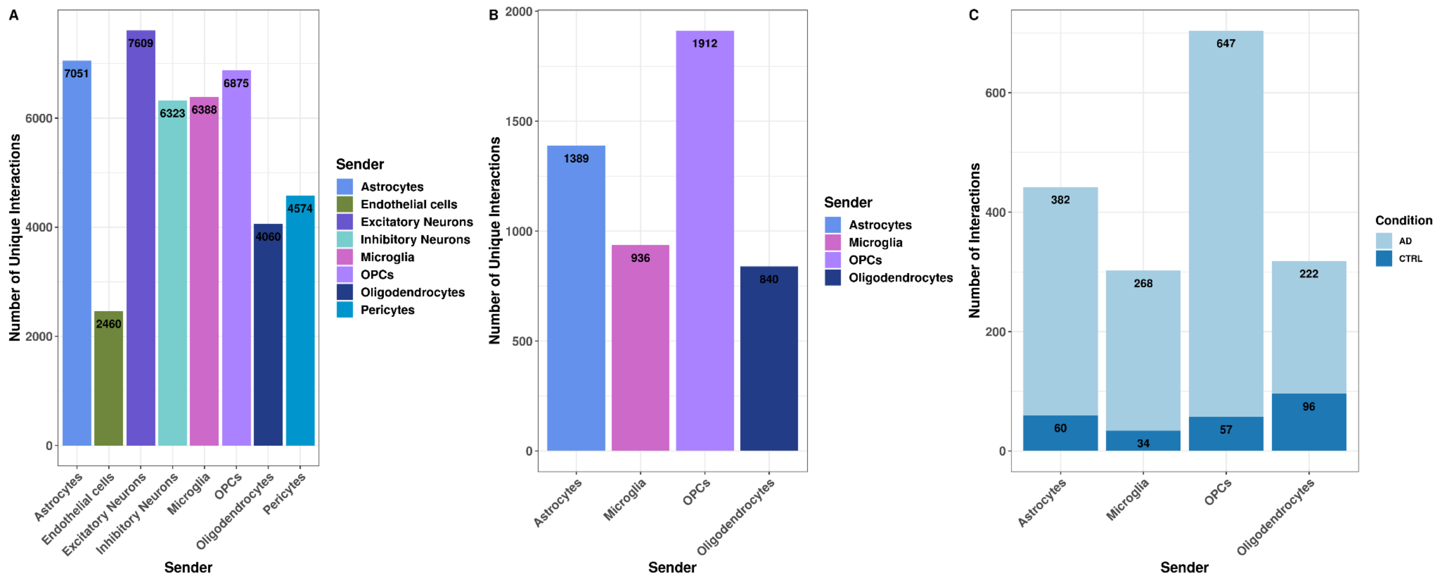

**Figure S4: Lau et al. cell type assignment and proportions**

(**A**) Uniform Manifold Approximation and Projection (UMAP) of 32,003 nuclei colored by their assigned cell types. (**B**) Violin plot of canonical marker gene expression used for cell type assignment. Color represents cell type. (**C**) UMAP of integrated dataset split by condition (AD, CTRL). (**D**) Stacked barplot of cell type proportions across conditions (AD, CTRL).

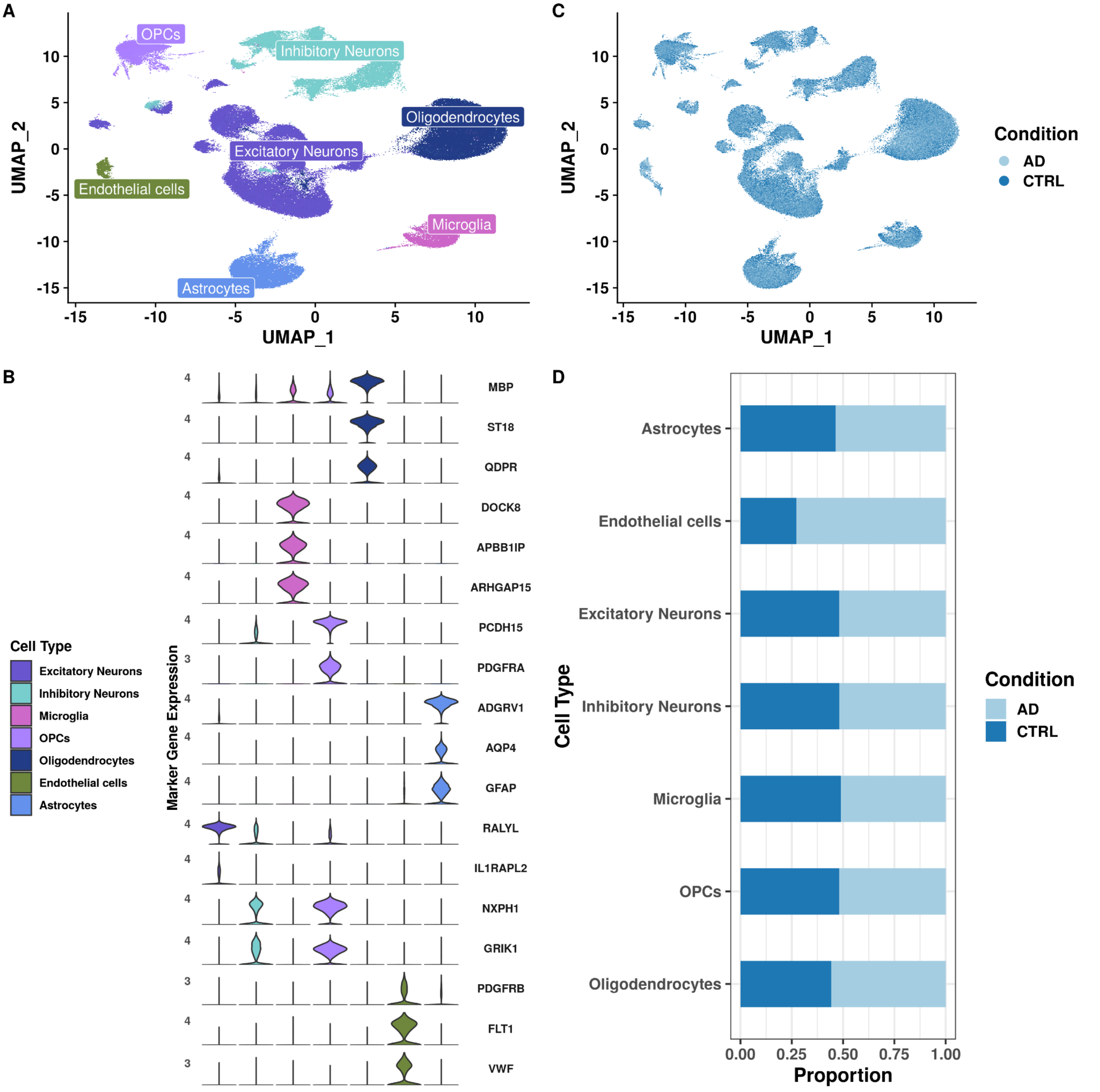

**Table S1: Morabito et al. Marker Genes**
Table of marker genes in GSE174367. All genes are upregulated in their respective clusters and have an average log2fc > 0.2. We used the top 25 markers based on their adjusted p-value for each cluster for cell type assignment. Table_S1.xlsx

**Table S2: Lau et al. Marker Genes**
Table of marker genes in GSE157827. All genes are upregulated in their respective clusters and have an average log2fc > 0.2. We used the top 25 markers based on their adjusted p-value for each cluster for cell type assignment. Table_S2.xls

**Table S3: Canonical marker genes**
Table of canonical marker genes used for cell type assignment using feature plots.

| **Canonical cell type markers** | **Cell type** |
|---|---|
| *PDGFRB* | Pericytes |
| *DOCK8*, *APBB1IP*, *ARHGAP15* | Microglia |
| *SNAP25*, *RBFOX1* | Neurons |
| *AQP4*, *GFAP*, *ADGRV1* | Astrocytes |
| *ST18*, *MBP*, *QDPR* | Oligodendrocytes |
| *PDGFRA*, *PCDH15* | OPCs |
| *FLT1*, *VWF* | Endothelial cells |

**Figure S5: Differential CCC analyses show low similarity of ligands, receptors, and targets across sender and receiver cell populations in Lau et al.**

**(A)** Stacked barplot of the number of prioritized interactions where excitatory or inhibitory neurons are the receiver. **(B)** Jaccard Similarity Index (JI) between excitatory and inhibitory neurons of receptors and targets. JI between sender cell type **(C)** ligands, **(D)** receptors, and **(E)** targets for inhibitory neurons**.** JI between sender cell type **(F)** ligands, **(G)** receptors, **(H)** targets for excitatory neurons.

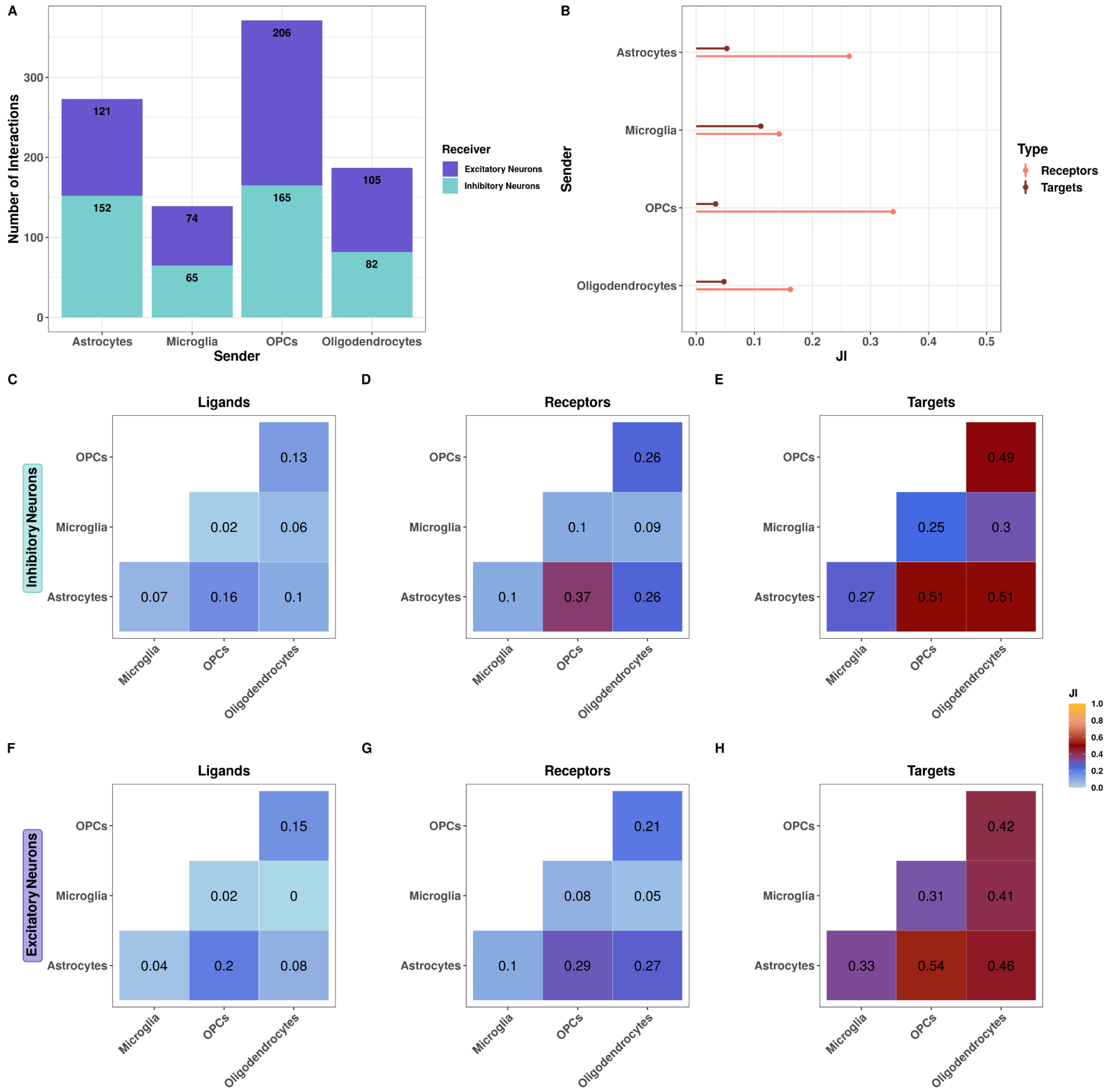

**Figure S6: Sadick et al. cell type assignment and proportions**

(**A**) Uniform Manifold Approximation and Projection (UMAP) of over 100,000 nuclei colored by their assigned cell types. (**B**) Violin plot of canonical marker gene expression used for cell type assignment. Color represents cell type. (**C**) UMAP of integrated dataset split by condition (AD, CTRL). (**D**) Stacked barplot of cell type proportions across conditions (AD, CTRL).

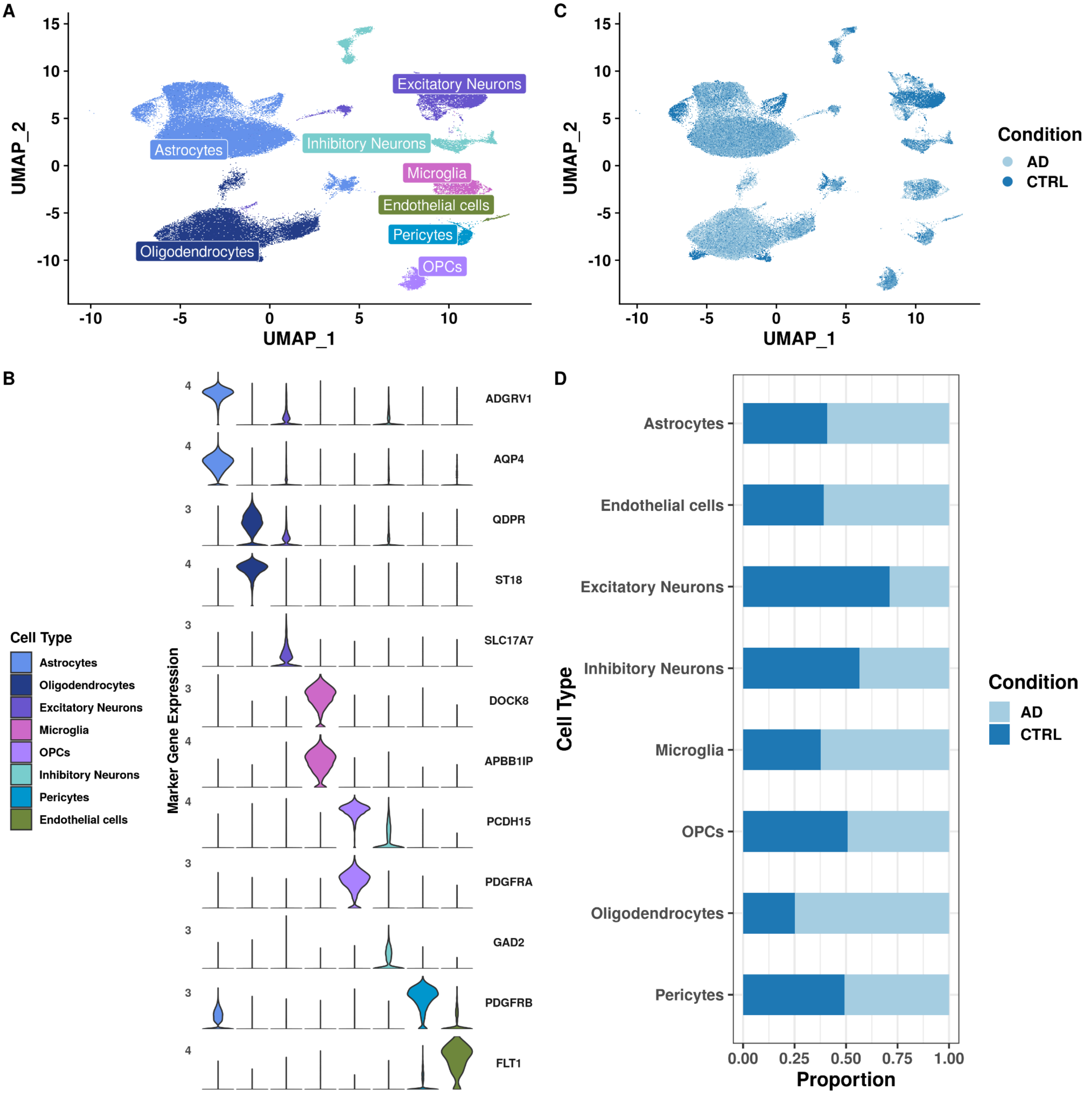

**Figure S7: Ligand-receptor and LRT overlap across all three snRNA-seq datasets**

(**A**) Upset plot of the overlapping ligand-receptor pairs **(B)** and LRTs. **(C)** Alluvial plot of the ligand-receptor pairs shared by all three datasets colored by sender cell type.

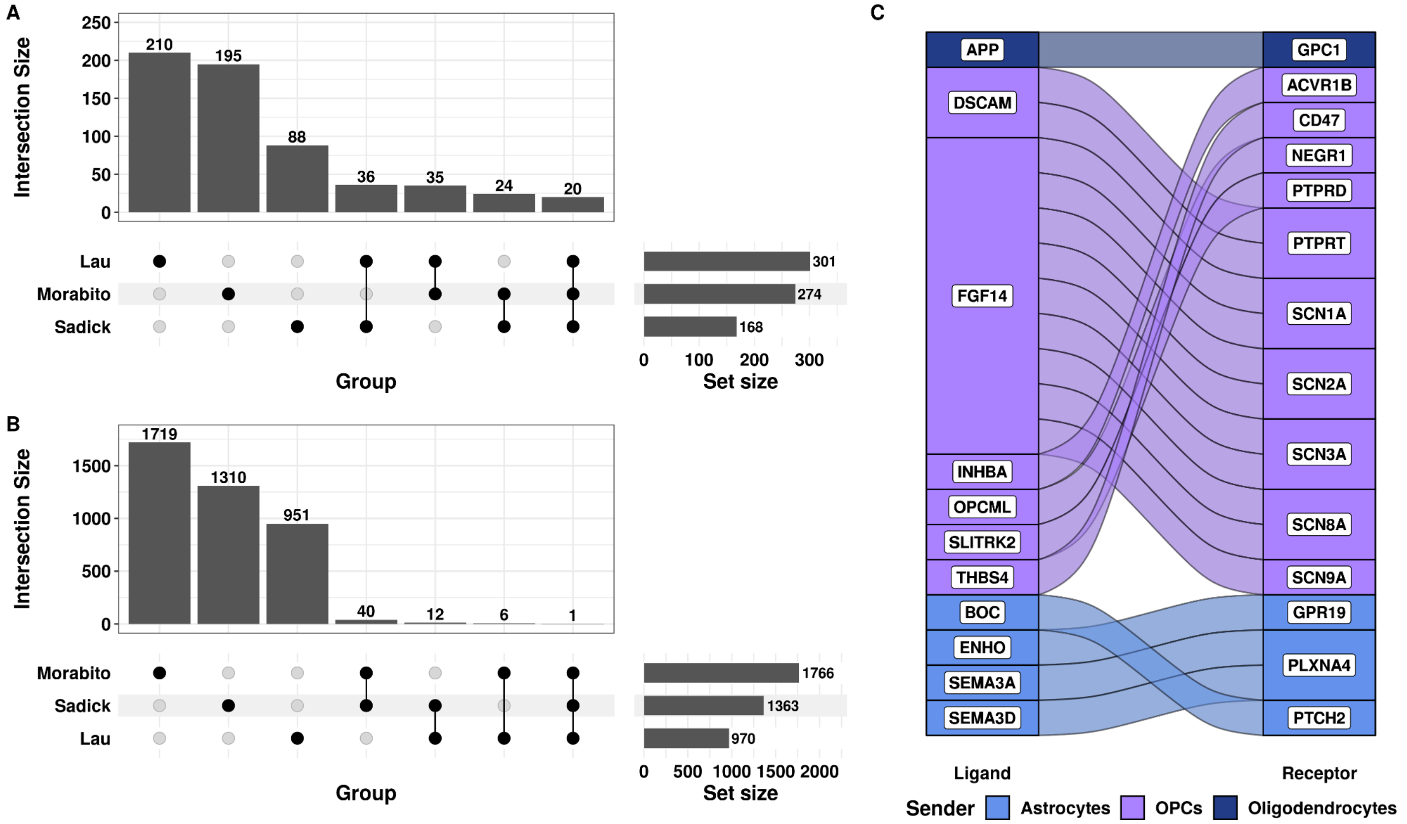

**Figure S8: Ligand-receptor pairs that overlap across two independent datasets. (A)** Overlapping ligand-receptor pairs that were upregulated in AD. **(B)** Overlapping ligand-receptor pairs that were upregulated in control. Colored by sender cell type. Grey indicates the involvement of more than one sender.

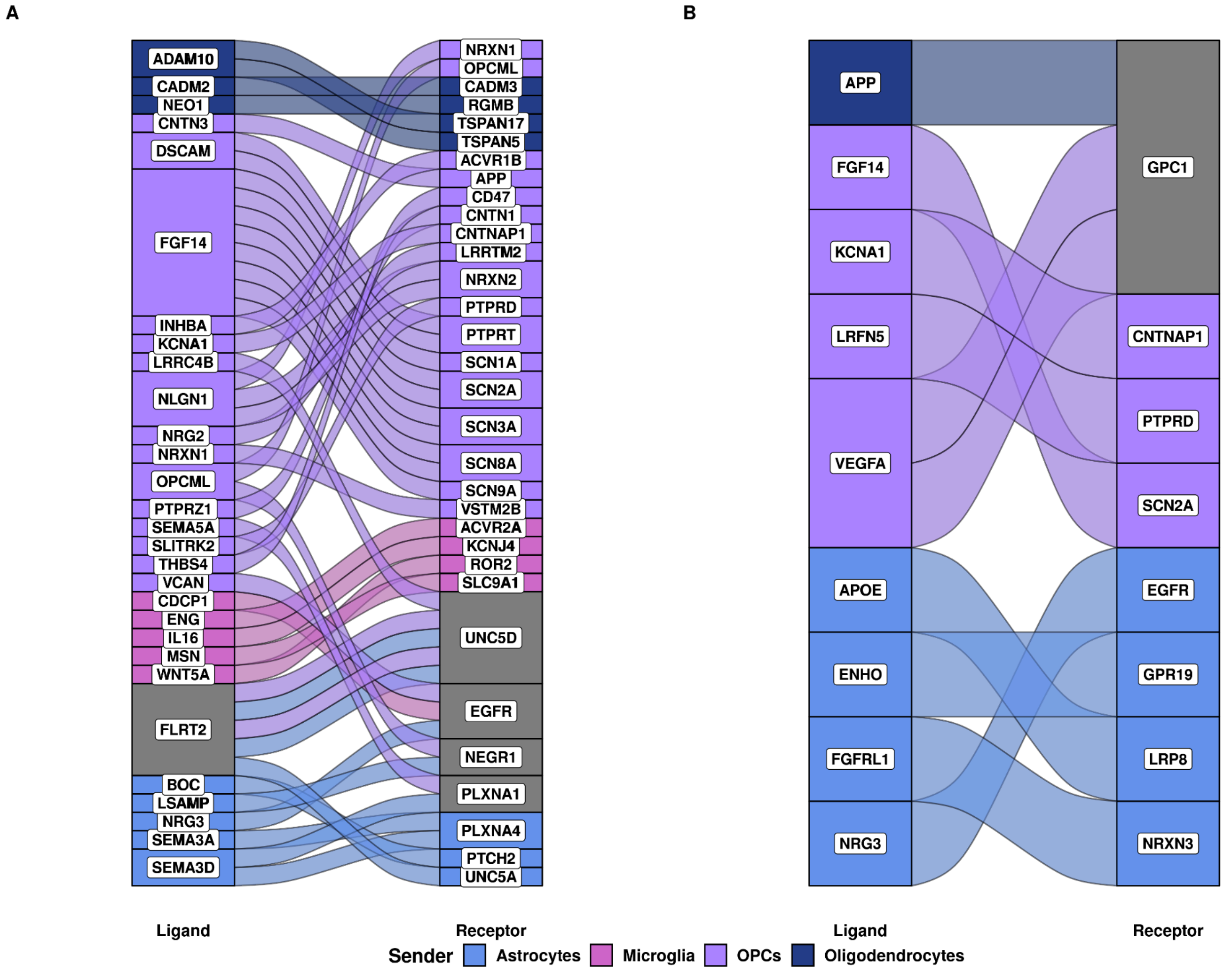


### Table S4: Target genes of ligand-receptor pairs that overlap across datasets

Table of 51 ligand-receptor pairs that overlap across datasets. The table includes the ligand-receptor pair, the sender and receiver cell type associated with the pair, as well as the target genes annotated by dataset of origin. Table_S4.xlsx

**Figure S9: Alluvial plot of high-confidence LRTs between Morabito and Lau before using patient sex as a covariate.**

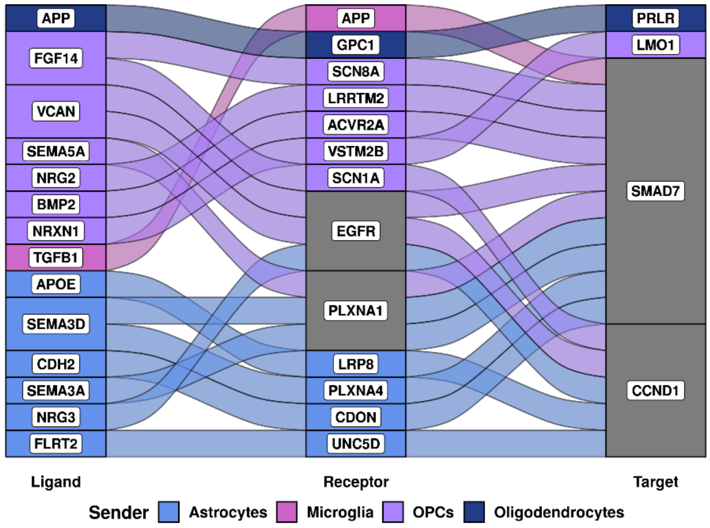
